# Embryonic cortical extracellular vesicles confer neuroprotection via multipathway signaling with CaMKIIα as a key mediator

**DOI:** 10.64898/2026.02.18.706575

**Authors:** Raquel García-Rodríguez, Sandra González de la Fuente, Marta Guerrero-Valero, Marta Carús-Cadavieco, Irene Clares Pedrero, Carlos Cabañas, Ernest Palomer, Francesc X. Guix, Carlos G. Dotti

## Abstract

Extracellular vesicles (EVs) are increasingly recognized for their roles in orchestrating embryonic development. Emerging preclinical evidence further suggests that EVs from young organisms possess innate regenerative potential for adult or injured tissues. Here we show that small extracellular vesicles (sEVs) isolated from the mouse embryonic cortex exert neuroprotective effects *in vitro* and *in vivo*. Proteomic profiling revealed that embryonic sEVs are enriched with effectors of receptor tyrosine kinase activation, anti-inflammatory responses, and protein synthesis. Notably, we identified BDNF as a surface-bound cargo on embryonic sEVs, displaying superior stability and receptor activation kinetics than its non-vesicular form. Phospho-proteomic analysis further revealed that sEV-mediated neuroprotection is driven primarily by the CaMKIIα signaling axis, which targets downstream effectors of microtubule stability, synaptic plasticity, and membrane-cytoskeleton interactions. Critically, embryonic sEVs, but not those from aged mice, restored microtubule stability and mitochondrial respiration in aged neurons in vitro. Our findings identify embryonic cortical sEVs as significant regulators of neuronal resilience and provide a molecular blueprint for EV-based strategies in neurodegeneration and aging research.

## Introduction

As post-mitotic cells, neurons remain in a long-term quiescent state throughout an organism’s lifetime. During aging, these cells accumulate structural, biochemical, and molecular deficits driven by a combination of cell-autonomous and non-autonomous mechanisms (Mattson & Arumugam, 2018). Although these impairments contribute to cognitive waning, extensive stereological studies have shown that significant neuronal attrition is not an inevitable feature of aging (Freeman et al., 2008; Morrison & Hof, 1997). This implies that neurons activate specific survival mechanisms to endure age-related stress. Nevertheless, the eventual breakdown of these homeostatic responses renders aging the leading risk factor for Alzheimer’s disease and other neurodegenerative conditions (Hou et al., 2019). Consequently, therapeutic strategies designed to augment these latent survival pathways represent a promising frontier for mitigating age-related dysfunction and forestalling the onset of neurodegeneration.

Various therapeutic strategies are currently being investigated to promote cellular longevity and allay age-related physiological decline. These include the activation of pro-survival signaling cascades, such as the PI3K/Akt and MAPK/Erk pathways (Beker et al., 2019; Hou et al., 2018), and the use of neurotrophic factor mimetics (Akhtar et al., 2021; Massa et al., 2010). Other molecular approaches focus on reducing chronic neuroinflammation (Costagliola et al., 2022; Ifergan & Miller, 2020), and enhancing autophagic flux to maintain proteostasis (Berger et al., 2006; Yang et al., 2014). Additionally, systemic interventions are being explored, ranging from the administration of young-blood-derived factors (Iram et al., 2022; Villeda et al., 2014) to lifestyle-based modulations of metabolic and cognitive health (Mirescu et al., 2006; Sleiman et al., 2016).

Beyond these strategies, extracellular vesicles (EVs) derived from young or embryonic tissues are emerging as potent mediators of systemic rejuvenation and tissue repair (reviewed in Becker et al., 2016; Hill, 2019; van Niel et al., 2022). As heterogeneous, membrane-bound vesicles secreted by virtually all cell types, EVs transport a diverse repertoire of bioactive molecules that modulate signaling pathways in recipient cells. Current nomenclature, standardized by the International Society for Extracellular Vesicles (ISEV), classifies these vesicles primarily by size: small EVs (sEVs), typically < 200 nm, and large EVs, which span from 200 nm to 10 µm (Théry et al., 2018). Exosomes represent a distinct sEV population defined by their endosomal origin. They are generated *via* the inward budding of late endosomal membranes, forming intraluminal vesicles (ILVs) within multivesicular bodies (MVBs). These MVBs follow one of two primary fates: fusion with lysosomes for cargo degradation or fusion with the plasma membrane to release ILVs into the extracellular environment as exosomes (Raposo & Stoorvogel, 2013). ILVs contain material from endocytosis, as well as other proteins, lipids, and nucleic acids from the cell cytosol (Dixson et al., 2023).

Functionally, EVs act as “double-edged swords”, either ameliorating or exacerbating pathological conditions depending on the cell type, developmental stage, and physiological state of both the donor and recipient cells. For instance, EVs secreted by senescent cells propagate inflammation and tissue dysfunction (Yin et al., 2021), while EVs derived from the blood-brain barrier have been implicated in the pathogenesis of multiple sclerosis (Dolcetti et al., 2020), and have been shown to drive cognitive deficits in Alzheimer’s disease models (Vandendriessche et al., 2021). Conversely, specific EV populations facilitate tissue repair and regeneration across various pathologies (reviewed in Wang & Pan, 2023).

Under physiological conditions, particularly in the brain, EVs play essential homeostatic roles. Brain-derived neurotrophic factor (BDNF) has been shown to modulate the cargo of neuron-derived EVs by inducing the sorting of specific miRNAs (miR-132-5p, miR-218-5p, and miR-690), which subsequently enhance excitatory synapse formation in recipient hippocampal neurons (Antoniou et al., 2023). Furthermore, neuron-derived EVs can modulate glial health, inhibiting microglial apoptosis (Peng et al., 2021) and promoting dendritic spine density *via* the TrkB-dependent phosphorylation of Akt and ribosomal protein S6 (RPS6) (Solana-Balaguer et al., 2023).

These functional outcomes appear highly dependent on the maturation state of the source neurons. Previous work from our laboratory demonstrated that sEVs derived from neurons maintained *in vitro* for 3–4 weeks––exhibiting signs of cellular distress––transport potentially toxic content into the extracellular medium (Guix et al., 2021). By contrast, sEVs from younger, 2-week-old cultures carry cargo that attenuates astrocyte reactivity (Almansa et al., 2022).

Building on evidence that young systemic or cerebrospinal fluid can reverse hallmarks of aging (Iram et al., 2022; Villeda et al., 2014), the present study investigated whether sEVs derived from the early-developmental mouse cerebral cortex exert neuroprotective effects. Our findings demonstrate that these embryonic sEVs exert potent neuroprotective effects and provide a mechanistic framework for their action, identifying specific molecular signaling axes that drive these phenotypic reversals.

## Results

### Small extracellular vesicles from embryonic cortical cells mitigate cellular damage *in vitro* and *in vivo*

We investigated the neuroprotective and reparative potential of sEVs derived from the cerebral cortex of embryonic day 16 mice (E16 EVs). This developmental stage was selected for its enriched biochemical profile, which is inherently optimized for rapid growth, intercellular communication, and survival. The isolation protocol and subsequent characterization of these vesicles, confirming their size, morphology, and the presence of canonical markers (tetraspanin CD81 and flotillin), alongside the absence of the endoplasmic reticulum marker calnexin, are detailed in Supplementary Figure 1 (S1A-D).

To assess neuroprotective efficacy, we utilized primary cortical cells maintained for 21 days *in vitro* (21DIV) as a model of chronic *in vitro* stress accumulation. At this stage, the cultures exhibited significant distress, characterized by increased lactate dehydrogenase (LDH) release (Fig. 1A), decreased activation of the anti-apoptotic kinase Akt (Fig. 1B), and elevated pro-inflammatory cytokine tumor necrosis factor-alpha (TNF-α) levels (Fig. 1C). For further evidence of stress accumulation with aging *in vitro*, see Martin et al., 2008; Martin et al., 2014; Martín-Segura et al., 2019 and Sodero et al., 2011. Incubation of the 21DIV cortical cells with E16 EVs significantly attenuated these markers of distress, resulting in decreased extracellular LDH (Fig. 1D), and increased levels of phosphorylated Akt (Fig. 1E). Crucially, these restorative effects were age-dependent; sEVs derived from 20-month-old mice (20m EVs) failed to elicit similar protective responses (Fig. 1 D-E). Fluorescent labeling with Bodipy confirmed that these sEVs effectively associate with and are internalized by the target cells (S1E).

**Figure 1.**
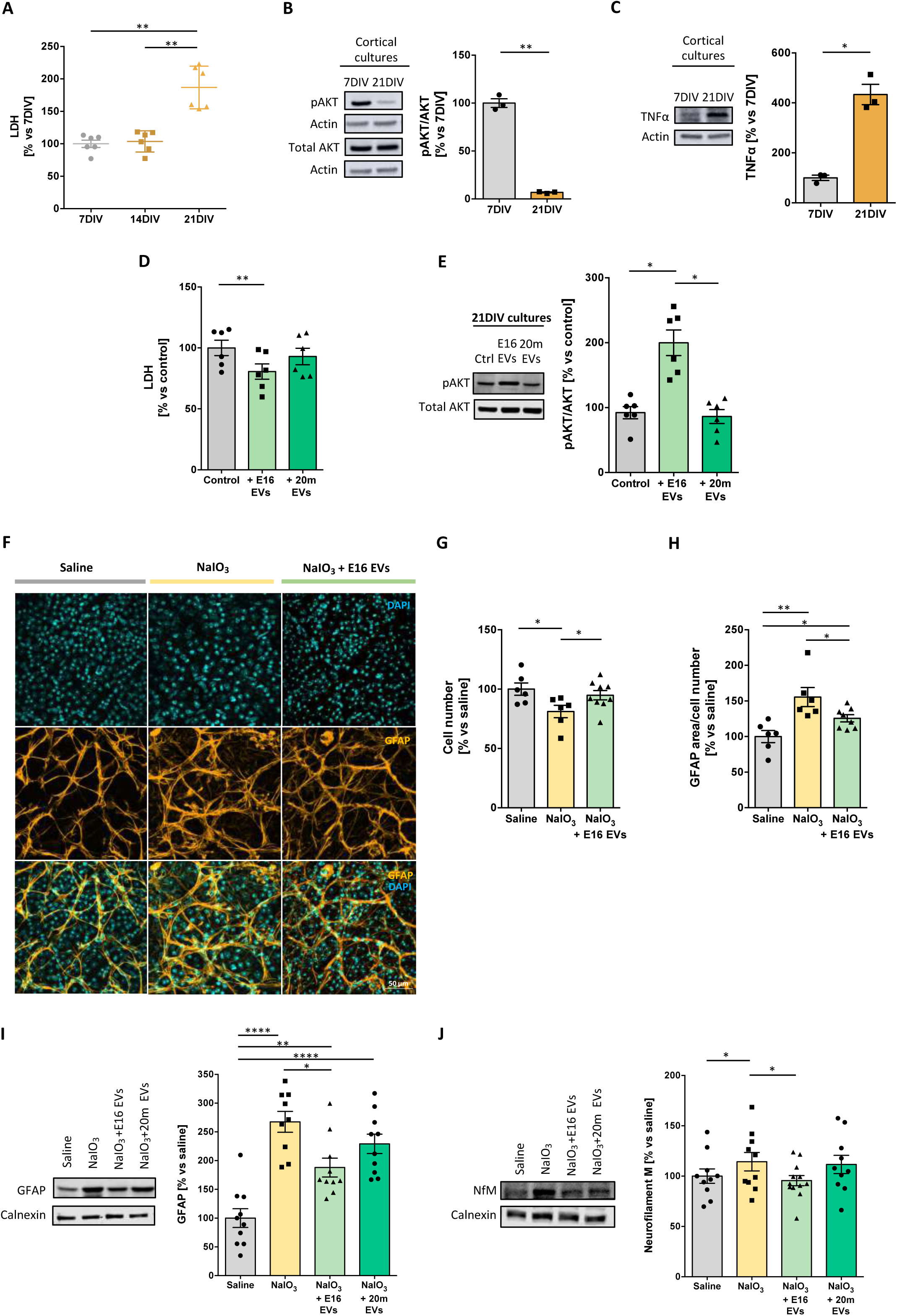
Small extracellular vesicles derived from mouse embryonic cortical tissue mitigate cellular stress in aged cortical neurons and attenuate inflammation in retinal degeneration. Primary cortical cells maintained for 21 days in vitro (21 DIV) were treated for 24 h with extracellular vesicles derived from the cerebral cortex of embryonic day 16 mice (E16 EVs) or 20-month-old mice (20m EVs) at 1 x 10^9^ particles/mL. All data represent the mean ± SEM as a percentage of the control (7DIV or untreated) values. **(A)** Absorbance measurement of LDH in conditioned medium at 7, 14, or 21DIV (n = 6; **p < 0.01; one-way ANOVA followed by Tukey’s post hoc correction). **(B and C)** Western blot analysis of the ratio of activated AKT (pAKT) to total protein (AKT) levels (B) and TNFα levels (C). Actin was used as a loading control (n = 3; *p < 0.05; Student’s t-test). **(D)** Absorbance measurement of LDH in conditioned medium of 21DIV cortical neurons treated or not with E16 EVs or 20m EVs (n = 6; *p < 0.01; one-way ANOVA followed by Tukey’s post hoc test). **(E)** Western blot analysis of the ratio of activated AKT (pAKT) to total AKT levels in 21DIV cortical cultures treated or not with E16 EVs or 20m EVs (n = 6; *p < 0.05; one-way ANOVA followed by Tukey’s post hoc test). **(F)** Representative retinal images from mice 5 days after intraperitoneal injection of 50 mg/kg NaIO_3_ followed by intraocular injection of PBS (NaIO_3_) or embryonic EVs (NaIO_3_+E16 EVs). Control animals received saline solution intraperitoneally and no further treatment (saline). Astrocytes are labeled with GFAP (yellow) and nuclei are counterstained with DAPI (blue). Data show the mean ± SEM expressed as a percentage relative to saline controls for cell number **(G)** and GFAP-positive area normalized to cell number **(H)**. n = 6/7; *p < 0.05 and **p < 0.01: one-way ANOVA followed by Tukey’s post hoc correction. **(I and J)** Representative western blot showing GFAP **(I)** or neurofilament-M **(J)** protein levels in mice subjected to intraperitoneal injection of 50 mg/kg NaIO₃ followed by intraocular treatment with E16 or 20m EVs. Data show mean ± SEM expressed as percentages relative to saline controls. Calnexin was used as loading control. n =9; *p < 0.05, **p < 0.01, and ****p < 0.0001; one-way ANOVA followed by Tukey’s post hoc correction.

Given the cellular heterogeneity of the E16 cerebral cortex—which comprises neuroblasts, radial glial cells, oligodendrocyte precursors, endothelial cells, and microglia—identifying the specific cellular origin of the neuroprotective sEVs *in vivo* is challenging. To isolate the potential contribution of the neuronal lineage, we established primary cortical cultures under serum-free conditions designed to selectively enrich for neuronal growth while suppressing the proliferation of non-neuronal populations. The isolation protocol, as well as the morphological and size characterization of these culture-derived sEVs, is detailed in Supplementary Figure 2 (S2A-C).

We evaluated the functional impact of sEVs derived from cortical neurons maintained *in vitro* for seven days (7DIV EVs). This timeframe was selected because 7DIV neurons reach a stage of significant morphological differentiation and metabolic activity, enabling robust intercellular communication and sufficient EV production for functional assays. The administration of 7DIV EVs to the stressed 21DIV cortical cultures decreased extracellular LDH levels (S2D), and increased the levels of phosphorylated Akt (S2E). Furthermore, 7DIV EVs exhibited potent neuroprotective capacity when cells were challenged with TNF-α (S2F). While these serum-free primary cultures are highly enriched for neurons, they may contain a residual population of astrocytes. Consequently, these results suggest that both neurons and astrocytes at early developmental stages serve as potential sources of EVs capable of exerting beneficial, restorative effects on stressed recipient cells.

To determine whether the neuroprotective capacity of embryonic sEVs extends to *in vivo* systems, we utilized a sodium iodate (NaIO_3_)-induced model of retinal degeneration (Enzbrenner et al., 2021). Five days following systemic NaIO_3_ administration, mice exhibited a significant reduction in total retinal cell count (Fig. 1F-G). This loss was accompanied by a significant increase in both glial fibrillar acidic protein (GFAP)-immunoreactive area and GFAP expression levels (Fig. 1H-I), signaling reactive gliosis in Müller cells and astrocytes. Additionally, we observed elevated levels of the neurodegeneration marker neurofilament M (NfM) (Fig. 1J). When administered concurrently with systemic NaIO_3_, intravitreal injection of E16 EVs —but not 20m EVs—significantly attenuated these degenerative markers, preserving cell numbers and reducing both GFAP-positive reactive gliosis and NfM accumulation (Fig. 1F-J).

These results demonstrate a protective effect of E16 EVs against stress-induced cellular damage *in vivo*. In contrast, 20-month EVs did not provide comparable protection, indicating an age-related decline in EV-mediated efficacy.

### Identification of pro-survival mediators in embryo vesicle-associated cortical sEVs: biochemical and functional characterization

To identify the molecular constituents underlying the neuroprotective effects of embryonic sEVs, we performed a proteomic analysis of E16 EVs using mass spectrometry. Consistent with the developmental profile of the donor tissue, the most abundant peptides identified were associated with trophic signaling, protein synthesis, protein polymerization, and anti-inflammatory pathways (Fig. 2A). Given that both 7DIV EVs and E16 EVs enhance Akt phosphorylation (Figs. 1E and S2E), we hypothesized that these vesicles transport ligands capable of activating Receptor Tyrosine Kinase (RTK)/Akt cascades. As a proof-of-concept, we focused on BDNF, a critical mediator of neuronal differentiation and survival that is expressed in the prenatal cerebral cortex and increases during postnatal development (Esvald et al., 2023). Western blot analysis of sEVs, derived from both embryonic cortical tissue and primary neuronal culture, confirmed the presence of the mature form of BDNF, and the absence of the precursor proBDNF (Fig. 2B). Notably, the levels of the mature form were higher in embryonic sEVs than in those isolated from 20 month-old mouse cortex (Fig. 2C).

**Figure 2.**
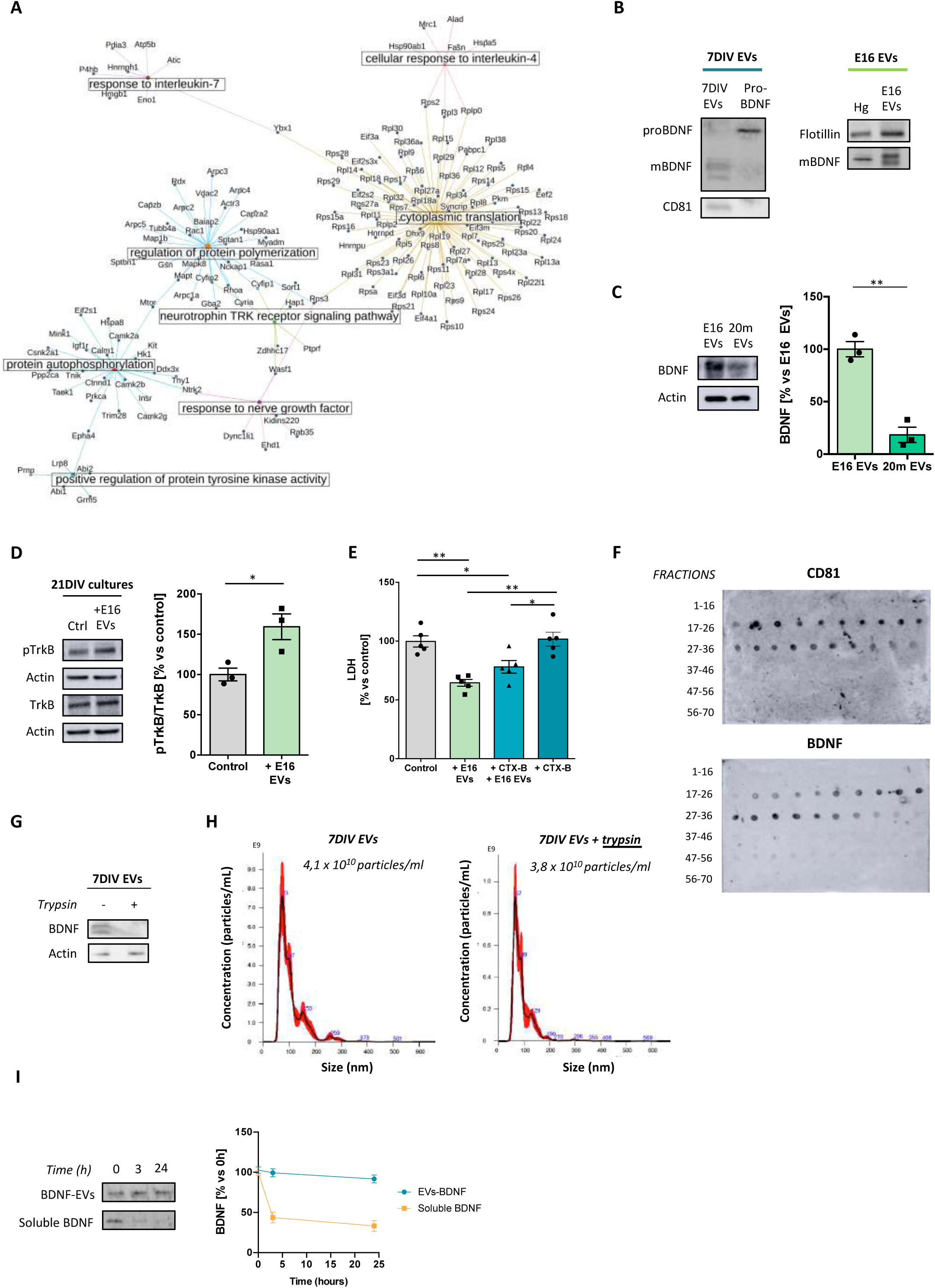
BDNF in embryonic sEVs is present on the outer surface, exhibits remarkable stability, and triggers TrkB activation in aged cortical neurons. **(A)** Pathway-based functional clusters according to peptide abundance of E16 EVs. Note that three clusters are functionally linked to receptor tyrosine kinase activity: neurotrophin TRK receptor signaling pathway, response to nerve growth factor, and positive regulation of protein tyrosine kinase activity. **(B)** Western blot confirming the presence of mature BDNF in both purified vesicles from cortical cells maintained for 7 days in vitro (7DIV EVs) and cortical cells from 16-day-old embryos (E16 EVs). The absence of the precursor form, pro-BDNF, is also shown in 7DIV EVs. Hg = homogenate **(C)** Western blot analysis showing lower levels of mature BDNF in purified vesicles from 20-month-old mice (20m EVs) compared with E16 EVs. Actin was used as loading control. Data represent the mean ± SEM as a percentage of E16 EV values. n = 3; *p < 0.05; Student’s t-test. **(D)** Western blot analysis of the ratio of activated TrkB (pTrkB) to total TrkB protein in 21DIV cortical cultures treated or not with E16 EVs (n = 3; *p < 0.05; Student’s t-test). Actin was used as a loading control. **(E)** Absorbance measurement of LDH in conditioned medium of 21DIV cortical cultures treated or not with cyclotraxin B (CTX-B, a selective TrkB inhibitor) and E16 EVs (n = 5; *p < 0.05 and **p < 0.01; two-way ANOVA followed by Tukey’s post hoc correction). Note that the addition of the TrkB inhibitor alone had no effect and only partially prevents the effect of the EVs. **(F)** Size exclusion chromatography of embryonic day 16 cerebral cortex extracellular vesicles (E16 EVs). Fractions were analyzed by dot blot and revealed the presence of BDNF exclusively in EV-rich fractions, as confirmed by the expression of the specific marker CD81. **(G)** Western blot analysis of the mature form of BDNF in 7DIV EVs, showing its loss after treatment of the EVs with 45 μg/mL trypsin for 30 min at 37 °C. **(H)** Particle size distribution and concentration measured by NTA before and after trypsin treatment. **(I)** Western blot analysis of the stability of BDNF present in 7DIV EVs compared with soluble BDNF. The presence of the mature form of BDNF was assessed at three different time points: 0, 3, and 24 h. Data show the mean ± SEM expressed as the percentage of BDNF levels normalized to the values measured at 0 hours. n = 3

To determine whether BDNF in sEVs is responsible for the observed neuroprotection, we quantified the activation of its specific receptor, TrkB, in recipient 21DIV neurons after EV incubation. Treatment with E16 EVs induced a significant increase in TrkB phosphorylation (Fig. 2D). Notably, the ability of E16 EVs to reduce LDH release was attenuated when recipient cells were pre-incubated with a pharmacological TrkB inhibitor (Cazorla et al., 2010), establishing that the neuroprotective effect is dependent on TrkB signaling (Fig. 2E). To further validate this mechanism, we utilized C6 glial cells, which primarily express the truncated TrkB.T1 isoform––lacking the intracellular kinase domain––rather than full-length pro-survival TrkB receptor (Ohira et al., 2006). In these cells, which exhibit reduced full-length TrkB expression compared to primary cortical neurons (S3A), E16 EVs failed to mitigate TNF-α-induced LDH release (S3B). While these results identify BDNF as a key functional component, the proteomic complexity of E16 EVs (Fig. 2A) suggests that the total neuroprotective effect likely results from the synergistic action of multiple trophic factors and signaling molecules transported by these vesicles.

A critical question arising from these findings is whether the detected BDNF is an intrinsic component of the sEVs or a soluble contaminant co-purified from the extracellular milieu. To distinguish between these possibilities, we fractionated the E16 EV preparations using size-exclusion chromatography. BDNF co-eluted with the canonical exosome marker CD81 (Fig. 2F), suggesting that BDNF is a *bona fide* vesicle-associated mediator rather than a co-isolated soluble protein. To determine whether EV-associated BDNF is sequestered within the lumen or tethered to the external surface, we subjected E16 EVs to trypsin-mediated proteolysis. Trypsin treatment effectively eliminated detectable BDNF (Fig. 2G), indicating that the neurotrophin is localized to the vesicle surface. To ensure that the loss was not due to protease-induced vesicle rupture, we utilized nanoparticle tracking analysis (NTA) to monitor biophysical integrity. Trypsin treatment resulted in no significant changes to particle size or concentration (Fig. 2H), confirming that the vesicles remained intact and that the BDNF pool is externally exposed. Finally, we investigated the temporal stability of EV-associated BDNF. While soluble BDNF typically exhibits a short half-life due to rapid enzymatic degradation (Mowla et al., 2001), the neuroprotective effects in our model persisted following overnight incubation. Comparative stability assays revealed that while soluble BDNF was degraded within 3 hours, EV-associated BDNF remained stable for at least 24 hours (Fig. 2I). These findings suggest that association with the EV membrane confers significant protection against proteolytic degradation, potentially extending the therapeutic window of the neurotrophin *in vivo*.

### Small embryonic EVs activate protective pathways in stressed neurons *in vitro*

To elucidate the molecular mechanisms underlying the protective effects, we performed a phospho-proteome mass spectrometry analysis on 21DIV cortical cells treated with E16 EVs. This analysis revealed widespread and significant shifts in the phosphorylation profiles of the target cells (Fig. 3A). Gene set enrichment analysis (GSEA) using the Gene Ontology (GO) database––specifically focusing on molecular function (MF), biological process (BP), and cellular component (CC)–-further characterized the functional implications of these changes (Fig. 3B). Among the multiple pathways affected, significant changes were observed in pathways governing cytoskeletal remodeling, neurogenesis, neuronal differentiation, and synaptic signaling. Given that E16 EVs significantly attenuate stress markers in 21DIV neurons (Fig. 1), these proteomic shifts likely represent a concerted pro-survival response. Supporting this, Kyoto Encyclopedia of Genes and Genomes (KEGG) pathway enrichment analysis confirmed that E16 EV treatment activates key signaling cascades essential for cell survival, such as the PI3K-AKT, MAPK, Ras, and mTOR pathways, alongside calcium-dependent signaling *via* CaMKII (Fig. 3C).

**Figure 3.**
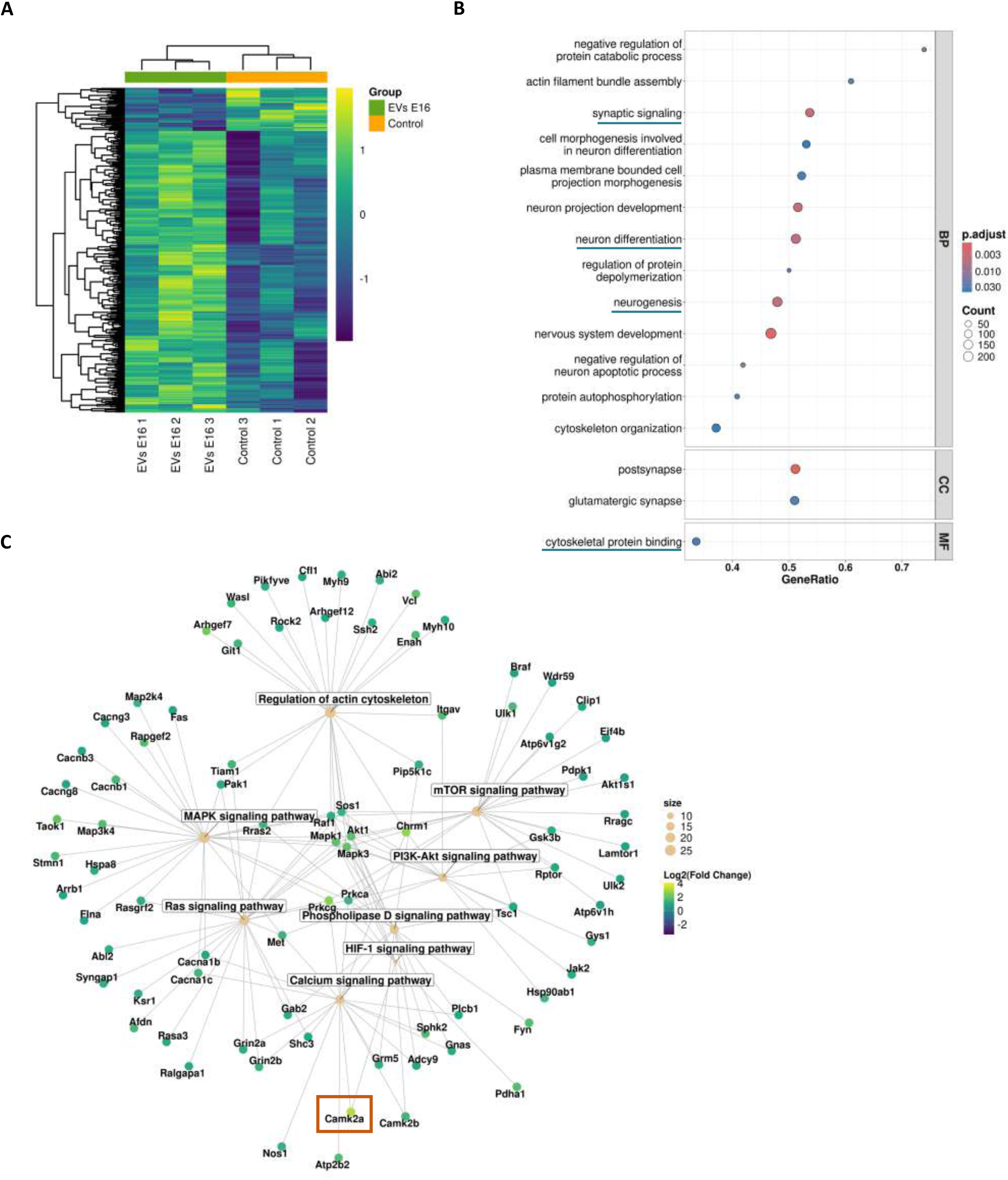
E16 EVs activate proteins involved in cell survival, neurogenesis, and synaptic plasticity in 21DIV cortical cultures. **(A)** Heatmap showing the expression profiles of significantly regulated phosphopeptides (p < 0.05) after treatment with E16 EVs. Each row represents a phosphopeptide and each column an individual replicate (n = 3 per group). Intensities are log2-transformed and row-scaled, with colors ranging from dark blue (downregulation) to yellow (upregulation). Group annotations (green: E16 EVs; orange: control) indicate sample condition. **(B)** Dot plot of enriched GO terms in treated cultures, grouped by GO domain: Cellular Component (CC), Molecular Function (MF), and Biological Process (BP). The x-axis represents the proportion of genes associated with each GO term and point color indicates the adjusted p-value. Dot size reflects gene count, with larger dots representing categories containing more genes. **(C)** Gene-concept network of selected KEGG pathways relevant to neuronal function in 21DIV neurons treated with EVs E16. Nodes represent phosphoproteins detected in the dataset, with edges connecting genes to enriched KEGG pathways. Node color indicates log2 fold-change, with a gradient from darkblue (downregulation) to yellow (upregulation), and node size corresponds to the pathway’s normalized enrichment score (NES).

### Small embryonic EVs promote CaMKIIα-dependent neuroprotection through differential modulation of downstream targets compared to aged EVs

Proteomic analysis identified the alpha isoform of CaMKII (CaMKIIα) among the top ten most significantly activated proteins in neurons treated with E16 EVs (Fig. 4A). This activation was confirmed by Western blotting of 21 DIV cortical cultures, which also showed greater CaMKIIα activity following E16 EV treatment than after treatment with 20m EVs (Fig. 4B). To establish a causal link between CaMKIIα activation and neuroprotection, we pharmacologically inhibited the kinase using KN93, which prevents activation by blocking calmodulin binding (Wong et al., 2019). While E16 EV treatment alone significantly reduced LDH release, the presence of KN93 abrogated this protective effect (Fig. 4C). Together, these data support a critical role for CaMKIIα in mediating the beneficial effects of sEV treatment.

**Figure 4.**
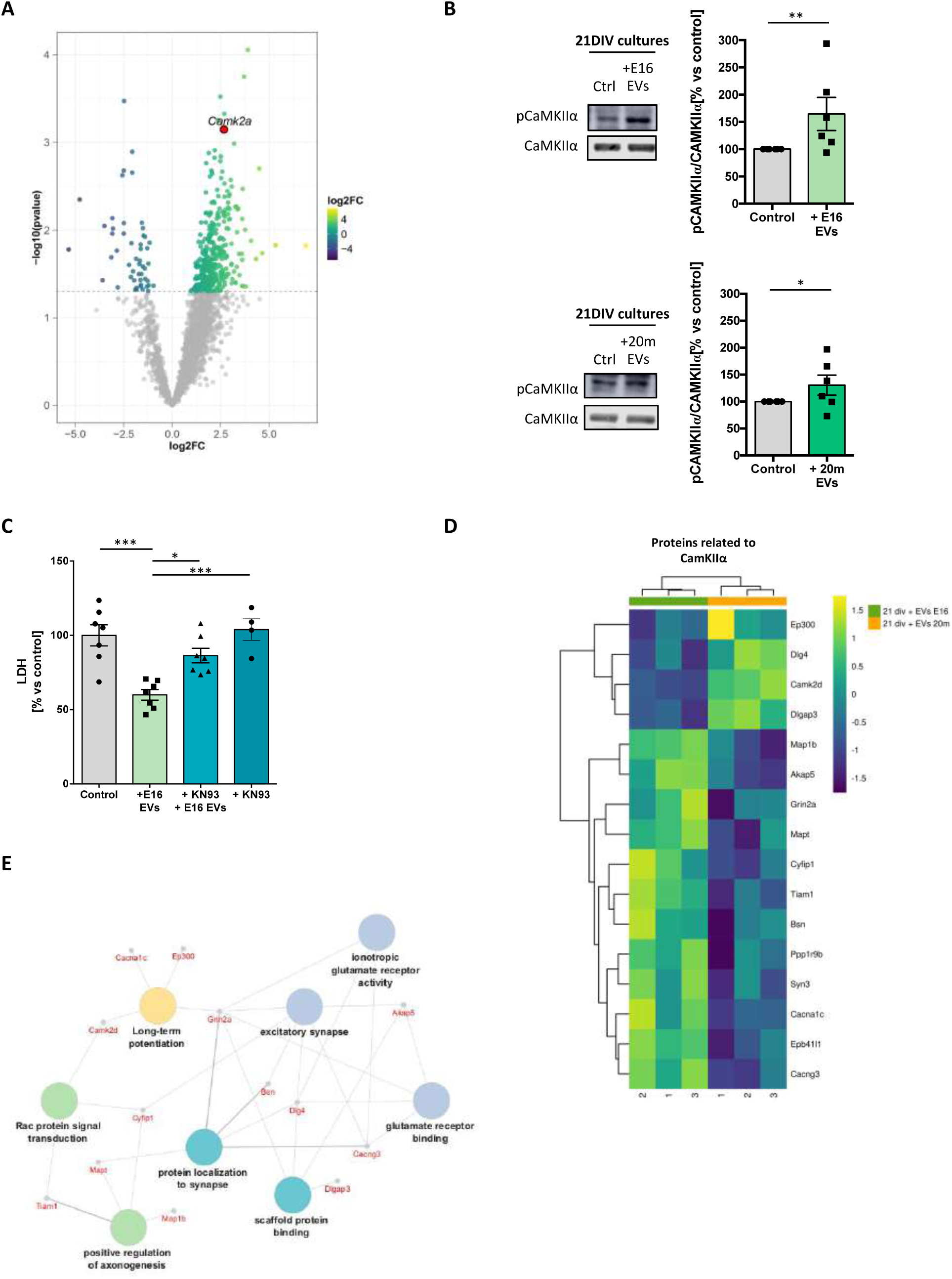
Embryonic sEVs induce CaMKII activation and selectively modulate CaMKIIα signaling cascades in 21DIV cortical cultures compared with aged sEVs. **(A)** Volcano plot showing the expression pattern of phosphopeptides in 21DIV cortical cultures treated with E16 EVs. The x-axis shows log2FoldChange and the y-axis shows −log10(p-value). Significantly up- or downregulated phosphopeptides are colored from yellow to darkblue while grey dots indicate peptides that were not differentially expressed. CAMK2a is highlighted in red with its gene name labeled. A dashed horizontal line represents the p-value cutoff (0.05). **(B)** Western blot analysis of the ratio of activated CaMKIIα (pCaMKIIα) to total CaMKIIα levels in 21DIV cultures treated or not with E16 or 20m EVs (n = 6; *p < 0.05; one-sample t-test). Data show mean ± SEM expressed as a percentage and normalized to N (control=100%). **(C)** Absorbance measurement of LDH release in the conditioned medium of 21DIV cultures treated or not with KN93 (selective CaMKII inhibitor) and E16 EVs. The addition of the KN93 inhibitor alone has no effect on LDH release. Data represent the mean ± SEM, expressed as a percentage and normalized to control values (n = 7; *p < 0.05 and ***p < 0.001; two-way ANOVA followed by Tukey’s post hoc correction). **(D)** Heatmap of the expression profiles of CaMKIIα effectors or related phosphopeptides in 21DIV cortical cultures treated with E16 EVs compared with those treated with 20m EVs. Each row represents a phosphopeptide and each column an individual replicate (n = 3 per group). Intensities are log2-transformed and scaled by row, with colors ranging from dark blue (downregulation) to yellow (upregulation). **(E)** Conceptual network of CaMKIIα effectors or associated peptides that are differentially expressed after treatment with E16 EVs and 20m EVs, and their relationship to biological processes.

Given that CaMKII activity is essential for EV-mediated neuroprotection, and that E16 EVs confer superior protection when compared with 20m EVs (Fig. 1), we hypothesized that embryonic sEVs activate a distinct set of downstream effectors. To test this, we performed phospho-proteomic analysis to identify CaMKII targets specifically modified by E16 EVs but not by 20m EVs. We identified sixteen such peptides: 12 were upregulated and 4 were downregulated (Fig. 4D). Bioinformatic clustering of the hyperphosphorylated targets revealed three primary neuroprotective mechanisms uniquely enriched by E16 EV treatment: (1) microtubule stability (Mapt/Tau, Map1B), (2) calcium and glutamate signaling (Cacng3, Cacna1, Grin2), and (3) membrane-synaptic cytoskeleton interactions (Akap5, Cyfip1, Bassoon, Synapsin-3, Tiam1, Epb4, Neurbin2/Ppp1R) (Fig. 4E).

### Small embryonic EVs enhance microtubule stability and maximal mitochondrial respiration

As sEVs carry complex cargo capable of engaging multiple neuroprotective pathways (Fig. 2A), their efficacy likely results from the activation of several targets rather than a single protein. To investigate this integrated response, we first assessed the impact of E16 EVs on microtubule stability. We focused on this mechanism because E16 EVs––unlike 20m EVs––specifically modulate Tau and MAP1B (Fig. 4D), proteins known to promote microtubule stability and protect against stress-induced depolymerization (Dubey et al., 2015). As shown by increased levels of tubulin acetylation (a marker of stable microtubules), E16 EV treatment, but not 20m EVs, enhanced microtubule stability in recipient neurons (Fig. 5A-B). Given the link between cytoskeletal integrity and metabolic health, we next evaluated mitochondrial respiration in 21 DIV cortical cultures treated with E16 or 20m EVs (Fig. 5C). While both E16 and 20m EVs induced a modest increase in basal respiration (Fig. 5D), only E16 EVs elicited a statistically significant enhancement in maximal mitochondrial respiration (Fig. 5E).

**Figure 5.**
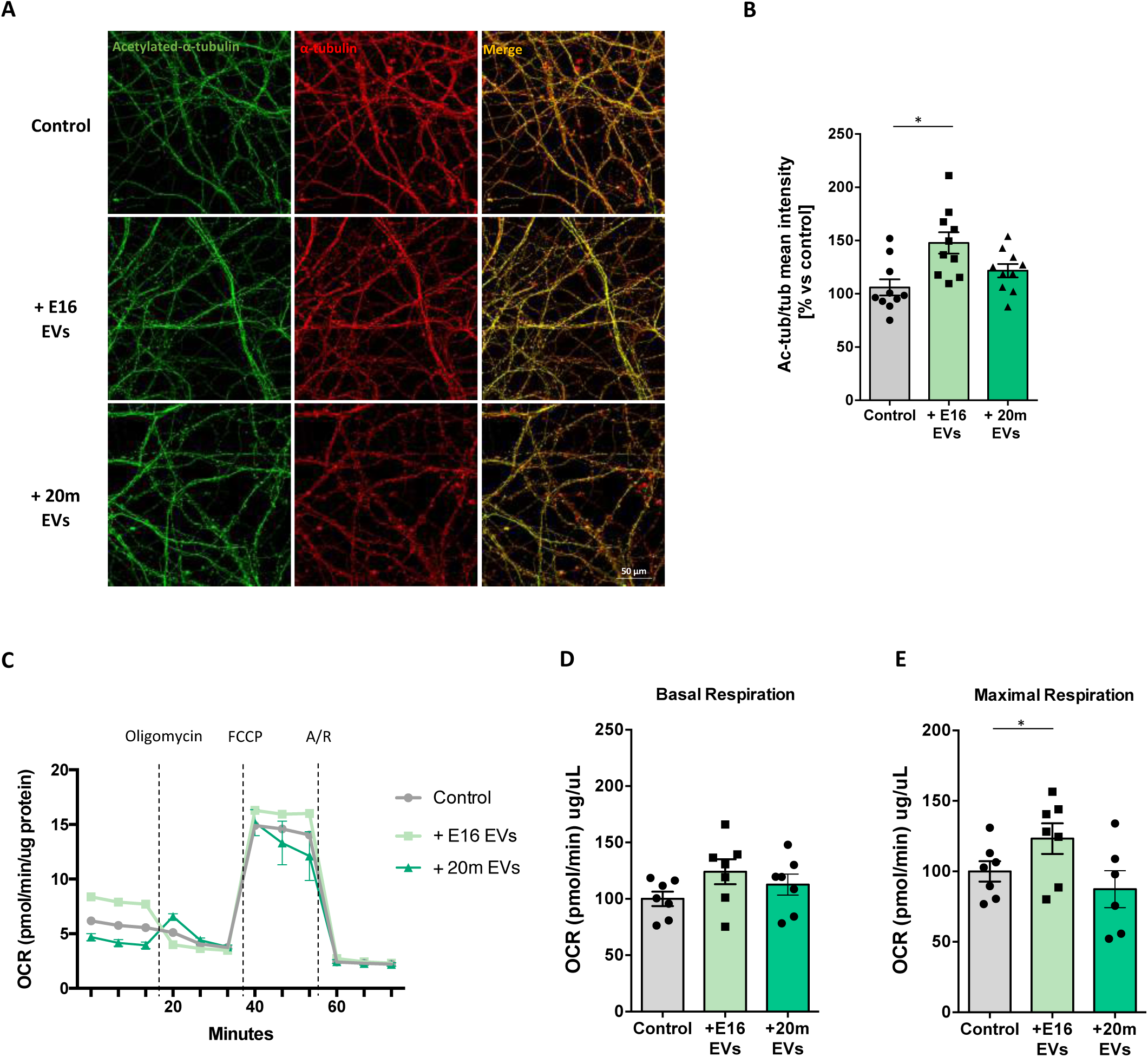
Small extracellular vesicles from embryonic cortex enhance microtubule stability and boost mitochondrial respiration in recipient cells. **(A)** Representative images of treatment with E16 EVs and 20m EVs in 21DIV cortical cultures at 1 x 10⁹ particles/mL for 24 h. Acetylated α-tubulin is shown in green and α-tubulin in red. Scale bar: 50 µm. **(B)** Fluorescence intensity corresponding to the acetylated α-tubulin/α-tubulin ratio is shown in 21 DIV cells treated or untreated with E16 or 20 m EVs. Data represent the mean ± SEM as a percentage relative to control values, with each point representing replicates from four independent cortical cultures (*p < 0.05, one-way ANOVA followed by Tukey’s post hoc test). **(C)** Representative Seahorse assay profile showing oxygen consumption rate (OCR) normalized to ug of protein, following sequential addition of oligomycin (ATP synthase inhibitor), FCCP (mitochondrial uncoupler), and antimycin/rotenone (A/R) (inhibitors of complexes III and I, respectively). Basal respiration **(D)** and maximal respiration **(E)** values are shown for 21DIV cultures treated or not with E16 and 20m EVs. Data represent the mean ± SEM, expressed as a percentage and normalized to control values (n = 6; *p < 0.05, one-way ANOVA followed by Tukey’s post hoc correction).

These results reveal two key mechanisms by which embryonic sEVs may exert their neuroprotective effects: increased microtubule stability and enhanced cellular metabolic efficiency.

## Discussion

A central contribution of this work is the demonstration that sEVs derived from the cerebral cortex of E16 mice significantly attenuate neural stress in both *in vitro* cortical cultures and an *in vivo* model of retinal degeneration. While the neuroprotective potential of neurostem-cell-derived EVs has been previously documented (Campero-Romero et al., 2023; Cossetti et al., 2014; Stronati et al., 2019), our study is unique in its focus on the heterogeneous EV populations of a critical developmental peak. At E16, the cortex is a complex milieu of neuroepithelial stem cells, radial glial progenitors, and intermediate progenitors, alongside nascent neurons and astrocytes (Levers et al., 2001; Sahara et al., 2020). Notably, our data using sEVs purified from primary cortical cultures suggest that vesicles secreted by differentiating neurons (and potentially astrocytes) are active contributors to this neuroprotective effect. By moving beyond purely phenotypic observations, our work provides a mechanistic framework that integrates EV molecular composition with the activation of specific pro-survival signaling cascades in recipient cells.

The neuroprotective potential of embryonic brain-derived EVs is particularly significant when contextualized within the broader field of systemic rejuvenation. Extensive research has already highlighted the anti-aging and disease-modifying benefits of “young” systemic environments, including plasma and other biological fluids (Katsimpardi et al., 2014; Sahu et al., 2021; Iram et al., 2022). However, utilizing EVs as the primary therapeutic vehicle offers several distinct advantages over whole fluids or tissues. First, EVs derived from early embryonic stages possess a much less complex molecular profile than fluids or tissue homogenates, which reduces the risk of adverse immune responses or undesired cell proliferation (El Andaloussi et al., 2013). Second, the modular nature of EVs allows for rigorous molecular characterization using standardized proteomic and transcriptomic techniques, enabling the identification of specific bioactive mediators. Crucially, EVs possess the intrinsic ability to cross the blood-brain barrier, a capability often lacking in the larger protein complexes or “beneficial” components present in systemic fluids, allowing them to exert effects directly within the central nervous system by delivering a specialized cargo of proteins, lipids, and nucleic acids from their cells of origin (Yáñez-Mó et al., 2015). Furthermore, EVs can be engineered to carry exogenous small-molecule compounds (Liu et al., 2023), significantly expanding their potential as a precision-medicine tool for treating brain pathologies.

A critical consideration in evaluating embryonic cortical sEVs is identifying the specific cell types that undergo recovery. Our findings suggest that E16 EVs exert protective effects on both neurons and astrocytes. The direct impact on neurons is demonstrated by the reduction in LDH release, the upregulation of AKT signaling, and the enhancement of mitochondrial bioenergetics in 21DIV cortical cultures. Simultaneously, the beneficial impact on astrocytes reactivity is evidenced *in vivo* by the marked reduction of GFAP expression in our retinal degeneration model. The robust repair observed in the retina further underscores the potential therapeutic versatility of E16 EVs in diverse neurodegenerative contexts. While our current study did not investigate the influence of these vesicles on microglia or oligodendrocytes, their broad efficacy across neurons and astrocytes suggests a versatile potential to mitigate damage across multiple CNS lineages. Future studies will be required to define the full spectrum of their cellular targets and systemic influence.

The molecular basis of the observed neuroprotection likely stems from the specialized cargo inherent to vesicles derived from developing brain cells––entities naturally primed for rapid division, migration, and intercellular signaling. Our proteomic analysis confirmed that E16 EVs are enriched with peptides associated with neurotrophin signaling, anti-inflammatory responses, and protein synthesis. While the presence of neurotrophins such as BDNF in neuronal EVs has been previously reported (Chung et al., 2020; Delgado-Peraza et al., 2023), a major finding of this study is the exceptional stability of EV-associated BDNF (Fig. 2I). This longevity is clinically significant, as the therapeutic application of exogenous BDNF has historically failed due to its extremely short half-life (Miranda-Lourenço et al., 2020). We hypothesize that the EV lipid bilayer and perivesicular carbohydrate coat act as a structural shield, protecting the neurotrophin from degradation. The functional relevance of this stabilized BDNF is clear; pharmacological blockade of the TrkB receptor significantly attenuated the neuroprotective effect, confirming that BDNF/TrkB signaling is a critical axis of the embryonic EV response. Intriguingly, while sEVs from aged brains (20m) also present BDNF on their surface, they fail to confer neuroprotection. This discrepancy suggests two potential scenarios: quantitative deficiency or qualitative divergence. sEVs from aged animals contain significantly lower levels of BDNF (Fig. 2C). This reduction may be driven by age-related alterations in membrane fluidity and stability (Martín & Dotti, 2022), which could impair BDNF retention on the EV surface. Alternatively, other components within embryonic sEVs might be necessary to elicit neuroprotection beyond neurotrophins, some of which may be absent in sEVs from older animals. Notably, our proteomic data identified a suite of peptides associated with anti-inflammatory, phosphorylation, and RNA-translation pathways present (almost) exclusively in embryonic sEVs. In this scenario, the neuroprotective effect of embryonic sEVs may rely on specific components. Further research is needed to identify which other elements of embryonic sEVs cooperate with neurotrophins—whether their effect is due to embryonic-specific components, a quantitative difference, or both.

Given the multi-modal composition of E16 EVs, it is logical that their application triggers a broad spectrum of signaling cascades. However, a striking and somewhat unexpected finding was the predominance of the CaMKIIα pathway, as it is not typically the primary pathway downstream of RTK receptors such as those activated by BDNF. While embryonic sEVs did activate the canonical BDNF/RTK-mediated PI3K/Akt pathway, the number of activated CaMKIIα-related peptides was notably higher. Conceptually, CaMKIIα likely drives neuroprotection through several convergent mechanisms. CaMKIIα can activate transcription factors such as CREB, promoting the expression of anti-apoptotic genes including Bcl-2 and Bcl-xL (Meller et al., 2005). Additionally, by modulating the dynamics of microtubule- and actin-associated proteins, CaMKIIα stabilizes dendritic architecture and synaptic spines (Lemieux et al., 2012). Our data showing enhanced microtubule stability following E16 EV treatment supports this. Because microtubule integrity is essential for axonal transport, intracellular signaling, and overall neuronal structure, this regulatory role directly contributes to neuronal survival (Alberti et al., 2022). Finally, emerging evidence indicates that CaMKIIα maintains mitochondrial integrity and ATP production (Andersen et al., 2024), consistent with our observation of enhanced maximal respiration. A key takeaway from our study is the likely synergy between these effects. Microtubule stability and mitochondrial function are bidirectionally linked: microtubules facilitate the transport and distribution of mitochondria, while mitochondrial ATP provides the energy required for cytoskeletal remodeling (Cho et al., 2021; Gu et al., 2022; Yadav et al., 2022).

An intriguing finding of our study is that CaMKII is activated following treatment with both embryonic (E16) and aged (20m) EVs. This suggests that neuroprotection is not a product of CaMKII activation *per se*, but rather the recruitment of a specific subset of downstream targets unique to the E16 EV response. Bioinformatics analysis identified 12 CaMKII effectors exclusively activated in neurons treated with embryonic sEVs. These targets are primarily involved in microtubule stability, calcium regulation, glutamate transmission, and plasma membrane-cytoskeleton interactions. The functional relevance of this specificity is underscored by our observation that E16 EVs––but not aged EVs––successfully reduce LDH release, enhance microtubule stability and boost mitochondrial respiration in neurons maintained in culture for three weeks. It is highly probable that the selective manipulation of other E16-specific CaMKII targets could replicate these protective effects, representing a promising avenue for future therapeutic intervention.

Beyond its fundamental scientific contributions, we believe this work possesses significant translational potential. Demonstrating that embryonic cortical sEVs can mitigate neuronal stress *in vitro* and *in vivo* opens the door for their development as a disease-modifying therapy. While these vesicles are not envisioned as a “cure-all”, their ability to stabilize cellular architecture and bioenergetics suggests they could be valuable in slowing the progression of acute trauma or chronic neurodegenerative diseases. Further research remains essential to delineate the full therapeutic window of these vesicles.

## Methods

### Mouse primary cortical cultures

Primary brain cortex cultures were prepared from E16 mice according to established protocols (Dotti et al., 1988; Kaech & Banker, 2006). Briefly, cerebral cortices were dissected and collected in ice-cold Hanks’ Balanced Salt Solution (HBSS) supplemented with Ca^2+^ and Mg^2+^. The tissue underwent enzymatic digestion with 0.25% trypsin and 0.5 mg/mL DNase (#1128493200, Roche) for 15 min at 37 °C. Following three washes in HBSS, the tissue was mechanically dissociated in plating medium (Minimum Essential Medium, MEM, supplemented with 10% horse serum and 20% glucose). The resulting cell suspension was filtered through a 70-µm mesh and quantified using a Neubauer chamber. Cells were seeded onto poly-L-lysine-coated dishes (#P2636-1G, Sigma Aldrich) at densities of 15,000 cells/cm^2^ for immunofluorescence (0.5 mg/mL coating) or 30,000 cells/cm^2^ for biochemistry (0.1 mg/mL coating). Cultures were maintained in a humidified incubator at 37 °C with 5% CO_2_. After 3 h, the plating medium was replaced with equilibrated Neurobasal medium (#21103-049, GIBCO, Invitrogen) supplemented with B27 (#17504-044, GIBCO, Invitrogen) and GlutaMAX (#35050-061, GIBCO, Invitrogen). After 7 days *in vitro* (7DIV), the medium was gradually transitioned to MEM supplemented with N2 (1/3 of the medium per day for 3 consecutive days). Cortical neural cultures were maintained until 21DIV, a stage characterized by representative markers of cellular aging in this model (Sodero et al., 2011).

### Rat glioma C6 cell line cultures

Cells were cultured in Dulbecco’s Modified Eagle Medium (DMEM, Gibco) supplemented with 10% fetal bovine serum (FBS, Gibco), 100 U/mL penicillin, and 100 μg/mL streptomycin. Cultures were maintained at 37°C in a humified atmosphere containing 5% CO_2_. Cells were subcultured when they reached 80-90% confluence. For experiments, cells were seeded at a density of 1 x 10^5^ cells/mL in 6-well plates and allowed to adhere for 24 hours before treatment.

### Extracellular vesicles

#### Isolation of small extracellular vesicles from mouse cerebral cortex tissue

Small extracellular vesicles (sEVs) were isolated from mouse cerebral cortex tissue using a protocol adapted from Vella et al. (2017). Briefly, cerebral cortices were incubated in filtered Hibernate solution (#A12476-01, Gibco) supplemented with 75 U/mL type III collagenase (#LS004206, Worthington) at a ratio of 800 µL per 100 mg of tissue. The digestion was performed at 37 °C for a total of 20 min; during this period, samples were inverted every 2 min for the first 10 min, homogenized at 15 min, and incubated for a final 5 min. Following digestion, the suspensions was centrifuged at 300 × g for 5 min at 4 °C, and the resulting supernatant was subjected to sequential centrifugation at 2,000 × g for 10 min and 10,000 × g for 30 min, both at 4 °C. For purification, a discontinuous sucrose gradient was prepared in 14 × 95 mm polypropylene ultracentrifuge tubes (#331374, Beckman Coulter) by layering 1 mL of 2.5 M (F3), 1.2 mL of 1.3 M (F2), and 1.2 mL of 0.6 M (F1) sucrose solutions prepared in filtered PBS. The 10,000 × g supernatant was layered onto the gradient and the total volume was adjusted to 12.5 mL with filtered Hibernate medium. Samples were ultracentrifuged at 180,000 × g for 3 h at 4 °C (acceleration 4, deceleration 4) using an SW40 ultracentrifuge rotor (Beckman Coulter). Individual fractions were collected and washed in filtered PBS by ultracentrifugation at 100,000 × g for 1 h at 4 °C. The resulting pellets were resuspended in 100 µL of filtered PBS. From this volume, 5 µL was used for NTA, 10 µL for electron microscopy, and the remainder for downstream applications. For western blot analysis, sEVs were resuspended in RIPA buffer (50 mM Tris-HCl pH 7.4, 150 mM NaCl, 1% Triton X-100, 0.5% sodium deoxycholate, 0.1% SDS) instead of PBS.

#### Isolation of small extracellular vesicles from cell culture medium

To isolate sEVs from conditioned medium, cells were maintained for 7 DIV and seeded in p150 culture plates at 30,000 cells/cm². At 7DIV, the medium was harvested and subjected to two sequential centrifugations of 200 × g and 2,000 × g for 10 min each at 4 °C to remove cellular debris. The supernatant was ultracentrifuged at 10,000 × g for 30 min using 38-mL Ultra-Clear centrifuge tubes (#344058) in a swinging-bucket rotor (TST28.38, Beckman Coulter) to eliminate apoptotic bodies and microvesicles. The resulting supernatant was layered onto a 4-mL 30% sucrose cushion (#84100, Millipore) and ultracentrifuged at 100,000 × g for 2 h. The 6-mL fraction containing the vesicles was recovered and washed with sterile-filtered PBS by ultracentrifugation at 100,000 × g for 1 h. The final EV pellet was resuspended in 100 μL sterile-filtered PBS, with 5 μL allocated for NTA and the remainder was used for treatments. For western blot analysis, sEVs were resuspended in RIPA buffer instead of PBS.

#### Isolation of small extracellular vesicles by size-exclusion chromatography

sEVs were isolated from conditioned media by size-exclusion chromatography following a described protocol (Cardeñes et al., 2021). Briefly, the media were centrifuged at 300 × g for 5 min to remove cells and debris, followed by centrifugation at 10,000 × g for 40 min at 4 °C to pellet microvesicles. To isolate sEVs, the remaining supernatant was concentrated to 15 mL *via* tangential flow filtration using a Vivaflow 50 R membrane (Sartorius), followed by a second concentration step to 0.5 mL using a 100-kDa Amicon Ultra-15 centrifugal filter (Merck KGaA). The concentrate was loaded onto a 25 mL Sepharose 4 Fast-Flow resin column and eluted with filtered PBS. Seventy fractions were collected, and the protein profile was determined using a microBCA protein assay kit (#23235, ThermoFisher Scientific), measuring absorbance at 540 nm with a Tecan GENios multifunction microplate reader. EV-positive fractions were detected by dot blot: 2 µL of each fraction were loaded onto a nitrocellulose membrane (Pall Life Science), blocked with 2% bovine serum albumin (BSA, Sigma-Aldrich) and probed with an antibody against the exosome-specific marker CD81 overnight at 4 °C.

#### Transmission electron microscopy

Isolated EVs were placed on glow-discharged 200-mesh formvar/carbon-coated copper grids and stained with 2% uranyl acetate in water for 1.5 min at room temperature, followed by 3 washes in water. Grids were visualized using a JEOL JEM-1010 transmission electron microscope (JEOL, Tokyo, Japan), and digital images were acquired using a 4K × 4K F416 CMOS camera (TVIPS, Gauting, Germany).

#### Labeling of extracellular vesicles with BODIPY

To monitor cellular uptake, EVs were labelled with the lipophilic dye BODIPY 493/503 (#D3922, ThermoFisher). EVs were incubated with 1 µM BODIPY in PBS for 1 h at 37°C, and non-incorporated dye was removed using size-exclusion chromatography columns (MW 3000, ThermoFisher). Labelled EVs were then added to cell cultures for immunofluorescence analysis. As a negative control, PBS incubated under identical BODIPY-labeling conditions was processed in parallel.

#### Nanoparticle tracking analysis

NTA was performed using a NanoSight NS300 system (Malvern Panalytical, Malvern, UK) equipped with a high-sensitivity cMOS camera and a 488-nm laser. EV concentration and mean particle size were quantified using NTA software v2.3 (Malvern Panalytical). For each sample, three 60-s videos were recorded and analyzed.

#### Trypsin treatment of extracellular vesicles and stability assay

To determine the localization of BDNF, EVs isolated from 7DIV conditioned media were subjected to outer surface digestion assay with 45 µg/mL trypsin for 30 min at 37 °C. Following incubation, EVs were lysed in RIPA buffer and processed for western blot analysis. For BDNF stability assays, EVs isolated from 7DIV cultures and recombinant BDNF (50 ng/mL; #B3795, Sigma-Aldrich) were incubated for 0, 3, or 24 h at 37 °C in N2 culture medium.

#### Cell treatments

For functional assays, 21DIV neurons were treated for 24 h with EVs (1 × 10⁹ particles/mL) isolated either from embryonic cortex or from 7DIV neuronal cultures. To assess neuroprotection under inflammatory stress, 14DIV neurons or C6 glial cells were challenged with TNFα (50 ng/mL, #315-01A, PrepoTech) for 24 h, both in the presence and absence of EVs (1 × 10⁹ particles/mL). For signaling pathway inhibition experiments, 21DIV neurons were pre-incubated for 30 min with cyclotraxin B (CTX-B, TrkB inhibitor; 1 µM) or KN93 (CaMKIIα inhibitor; 20 µM) prior to 24-h incubation with embryonic EVs at 37 °C.

#### Lactate dehydrogenase assay

To quantify cell death 24-h post-treatment with EVs, 50 µL of conditioned media per sample was harvested and used to determine LDH levels with the Pierce LDH Cytotoxicity Assay Kit (88954, Thermo Scientific). The conversion of LDH to formazan was detected by measuring the absorbance at 490 nm using a FLUOstar OPTIMA microplate reader (BMG LABTECH).

#### Western blotting

Cells and EVs were lysed in RIPA buffer supplemented with phosphatase inhibitors (Phosphatase Inhibitor Cocktail Set III, Merck) and protease inhibitors. Lysates were sonicated (10 pulses, 100% amplitude, 0.8 cycles) using a Labsonic® sonicator (Sartorius) and cleared by centrifugation at 14,000 × g for 10 min at 4 °C. Protein concentration was determined using the BCA assay (#23225, Thermo Fisher Scientific). Samples were denatured at 90 °C for 5 min in reducing loading buffer (25 mM Tris-HCl pH 6.8, 1% SDS, 3.5% glycerol, 0.4% β-mercaptoethanol, 0.04% bromophenol blue). Proteins were resolved by SDS-PAGE and electroblotted to nitrocellulose membranes (330 mA, 1.5 h) using a Bio-Rad Mini-Protean system. Membranes were blocked in 2% BSA in TTBS (TBS + 0.1% Tween-20) for 30 min and incubated with primary antibodies overnight at 4 °C (Table 1). After washing, membranes were incubated with HRP- or fluorophore-conjugated secondary antibodies for 1 h. Protein bands were visualized using ECL (Thermo Fisher Scientific) for HRP or Odyssey M (LI-COR) for fluorescent antibodies. Band densitometry was quantified with FIJI/ImageJ software.

**Table 1.** List of antibodies used in western blotting (WB) and immunofluorescence (IF) analysis. Primary antibodies are shown in yellow and secondary antibodies in green.

| Target | Company | Reference | Host | Dilution |  |
| --- | --- | --- | --- | --- | --- |
|  |  |  |  | WB | IF |
| AKT | Cell Signaling | 9272S | Rabbit | 1:1000 | - |
| AKT | Cell Signaling | 2920 | Mouse | 1:1000 | - |
| p-AKT (S473) | Cell Signaling | 4060S | Rabbit | 1:1000 | - |
| β-actin | SIGMA | A5441 | Mouse | 1:10000 | - |
| BDNF | Alomone | ANT-010 | Rabbit | 1:500 | - |
| Calnexin | Abcam | Ab22595 | Rabbit | 1:10000 | - |
| CaMKIIα | Abcam | Ab52476 | Rabbit | 1:1000 | - |
| pCaMKIIα | Abcam | Ab171095 | Mouse | 1:1000 | - |
| CD81 | Santa Cruz | sc-166029 | Mouse | 1:500 | - |
| Flotillin | BD Bioscience | 610821 | Mouse | 1:500 | - |
| GFAP | Merck | MAB3402 | Mouse | 1:1000 | 1:2000 |
| MAP2 | Biolegend | 822501 | Chicken | - | 1:2000 |
| Neurofilament M | Biolegend | 841001 | Rabbit | 1:1000 | - |
| TNFα | Santa Cruz | sc-52746 | Mouse | 1:200 | - |
| TrkB | BD Bioscience | 610101 | Mouse | 1:1000 | - |
| pTrkB | Cell Signaling | 4621S | Rabbit | 1:1000 | - |
| α-tubulin | Sigma-Merck | T5168 | Mouse | - | 1:1000 |
| Acetylated-α-tubulin | Cell Signaling | 5335S | Rabbit | - | 1:1000 |
| IgG mouse-HRP | Dako | P0260 | Rabbit | 1:2500 | - |
| IgG rabbit-HRP | Dako | P0448 | Sheep | 1:2500 | - |
| IRDye rabbit-680RD | LI-COR | 926-68071 | Sheep | 1:10000 | - |
| IRDye mouse-800CW | LI-COR | 926-32210 | Sheep | 1:10000 | - |
| IgG rabbit-Alexa 488 | Thermofisher | A-11008 | Donkey | - | 1:500 |
| IgG mouse-Alexa 555 | Thermofisher | A-31570 | Donkey | - | 1:500 |
| IgY chicken-Alexa 647 | Thermofisher | A-21449 | Sheep | - | 1:500 |

#### Immunofluorescence analysis

Cells were fixed in 4% PFA for 10 min at room temperature followed by three washes with PBS. Cells were then permeabilized and blocked with 0.1% Triton X-100/2% BSA for 10 min. Primary antibodies were diluted in blocking solution and incubated overnight at 4 °C (Table 1). Following PBS washes, appropriate secondary antibodies were applied for 1 h at room temperature. Nuclei were counterstained with DAPI (1:5000). Coverslips were then mounted using ProLong medium. Images were acquired using a Zeiss LSM710 confocal microscope and analyzed with FIJI/ImageJ.

#### *In vivo* experiments

To induce retinopathy, 3–4-month-old mice of both sexes received a single intraperitoneal injection of sodium iodate (NaIO₃, 50 mg/kg) or saline (0.9% NaCl) as a vehicle control. NaIO₃ is a well-established model for macular degeneration, characterized by initial retinal pigment epithelium degeneration followed by secondary photoreceptor loss (Bhutto et al., 2018; Xiao et al., 2017). On the same day, mice were anesthetized with isoflurane and received a bilateral intravitreal injection of either E16 EVs or 20m EVs, according to established procedures (Nocera et al., 2024). Animals were allowed to recover and were maintained in their home cages for 5 d.

For histological analysis, mice were euthanized *via* CO₂ inhalation and underwent transcardial perfusion with 20 mL of 0.9% NaCl followed by 20 mL of 4% PFA. Eyes were enucleated, and the corneas were incised to facilitate fixative penetration during a 2-h incubation in 4% PFA at room temperature. Following three 15-min PBS washes, retinas were dissected, permeabilized in 0.5% Triton X-100/2% BSA for 1 h, and incubated with primary antibodies overnight at 4 °C. Following PBS washes, secondary antibodies were applied for 1 h, followed by DAPI counterstaining and mounting. Retinas were imaged using a Zeiss LSM710 system and analyzed with FIJI/ImageJ. For biochemical analysis, retinas were harvested immediately following 0.9% NaCl perfusion, and lysed in RIPA buffer for protein extraction.

#### Mitochondrial function analysis using Seahorse XF96

Neurons were seeded at 30,000 cells/well in Seahorse XF96 microplates and maintained for 21 DIV. For treatment groups, 21DIV cultures were incubated with E16 EVs or 20m EVs for 24 h. Prior to the assay, the culture medium was replaced with Seahorse XF Base Medium supplemented with pyruvate (1 mM), glutamine (2 mM), and glucose (10 mM), and equilibrated for 1 h at 37 °C without CO₂. Sequential injections included oligomycin (1 µM), FCCP (0.5 µM), and rotenone/antimycin A (0.5 µM each). Oxygen consumption rate was recorded in real time using Wave software and normalized to protein content. Parameters analyzed included basal respiration, ATP production, maximal respiration, spare respiratory capacity, and non-mitochondrial respiration.

#### Proteomics

Proteomic profiling was conducted at the CNIC Proteomics Facility (Madrid, Spain). Samples, including isolated E16 EVs and 21DIV neurons treated with E16 or 20m EVs were processed according to a published protocol (Núñez et al., 2024). Subsequent bioinformatics analysis was carried out by the Computational Analysis Service of the CBM utilizing artificial neural networks.

##### Data preprocessing and selection of significant proteins

Raw quantitative protein data from vesicle samples with adjusted p-values below 0.05 were selected as statistically significant. For proteins with duplicate identifiers, only the entry with the most significant p-value was retained to ensure a unique representation of each protein.

##### Protein–protein interaction analysis of CAMKIIα effectors

Interactions between CAMK2a and candidate protein effectors from extracellular vesicles (EVs) at embryonic day 16 (E16) and 20 months (20m) were retrieved from the STRING database (http://string-db.org/) with a minimum required interaction score of 0.40.

##### Pathway enrichment analysis

Gene Set Enrichment Analysis (GSEA) (Subramanian et al., 2005) was performed using the ClusterProfiler R package (Yu et al., 2012) with predefined gene sets from the Gene Ontology (GO) and KEGG databases, which provide information on gene functions and molecular interaction pathways. This analysis was used to identify gene sets showing statistically significant, concordant differences between experimental conditions. All peptides were mapped to their corresponding UniProt IDs, and the associated −log10(p-value)×SIGN values were used as input for the analysis. To report terms on enrichment tests, a hypergeometric distribution was used and *P*-values were corrected for multiple testing by the Benjamini–Hochberg algorithm. Gene enrichment analysis visualization was performed using the enrichplot (https://yulab-smu.top/biomedical-knowledge-mining-book/) and ggplot2 (https://ggplot2.tidyverse.org) R packages. The pheatmap package (Kolde, 2025) was used to visualize the expression values of the significant peptides.

### Statistical analysis

Statistical analyses were performed using GraphPad Prism 8.0 (GraphPad Software). Data are presented in all graphs as mean ± standard error of the mean (SEM). Prior to statistical testing, data normality was assessed using the Shapiro-Wilk test. For simple comparisons, Student’s t-test was applied to data with parametric distribution. For multiple comparisons, normally distributed data with one variable were analyzed using one-way ANOVA followed by Tukey’s post hoc correction, while datasets with more than one variable were analyzed using two-way ANOVA followed by Tukey’s post hoc test. A p-value < 0.05 was considered statistically significant. Statistical significance is indicated in the figures as follows: *p < 0.05; **p < 0.01; ***p < 0.001; ****p < 0.0001.

---

**Supplementary Figure 1.**
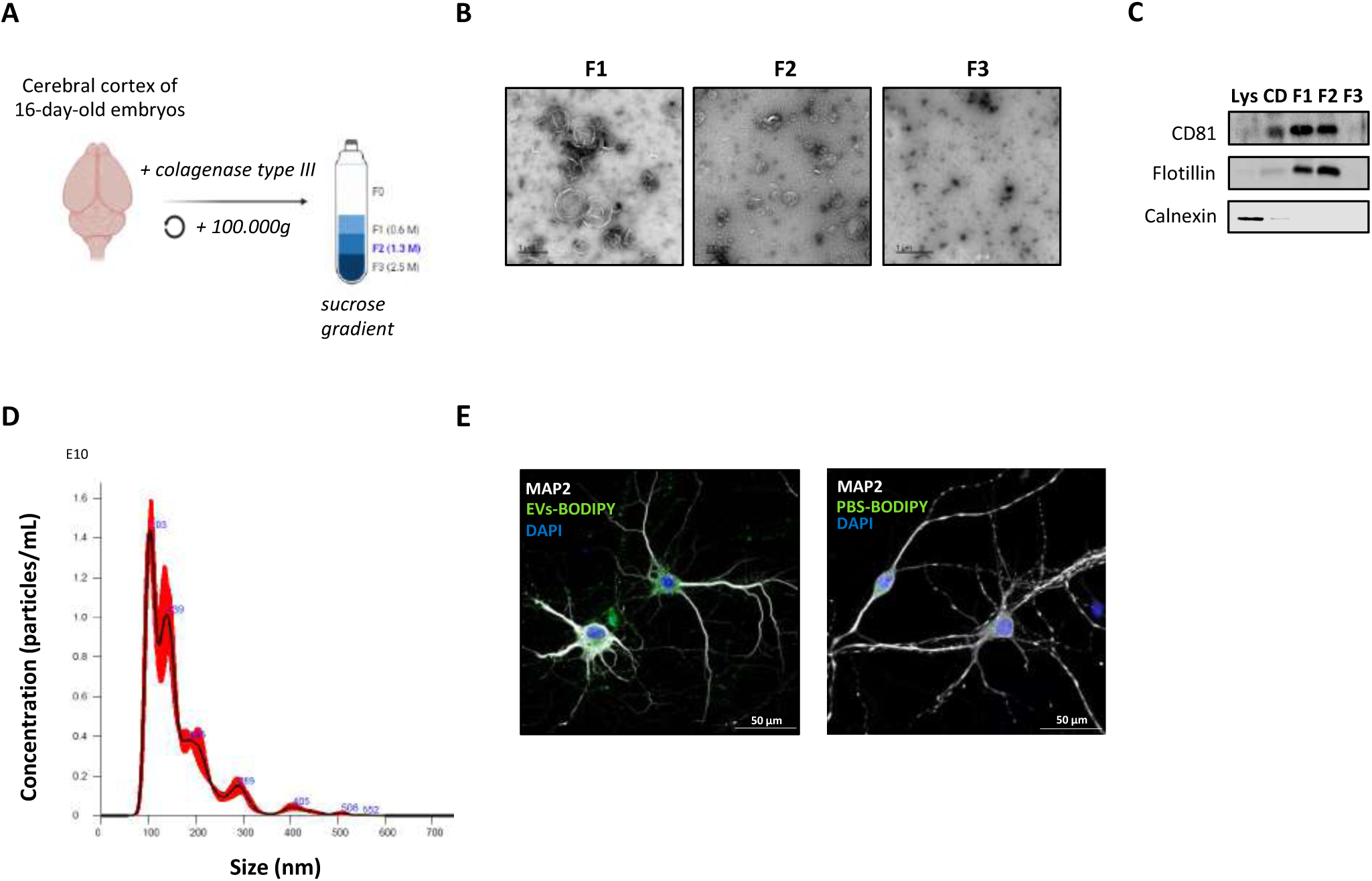
Isolation and characterization of small extracellular vesicles from mouse embryonic cerebral cortex. **(A)** Schematic representation of vesicle preparation from mouse embryonic cerebral cortex. **(B)** Representative electron microscopy images showing the different phases of the gradient used for vesicle isolation, with phase 2 (F2) being the most enriched in small-sized vesicles. **(C)** Western blot analysis of the different vesicle isolation phases, showing the enrichment of EV-specific markers such as CD81 and flotillin, and the absence of the endoplasmic reticulum marker calnexin in the fraction enriched in small vesicles (F2). **(D)** Representative plot showing the size distribution of the vesicles in phase 2 (F2) analyzed by Nanoparticle Tracking Analysis (NTA). Most vesicles have a size between 50 and 200 nm. **(E)** When extracellular vesicles labeled with the lipophilic dye Bodipy are incubated with cortical cultures, the neurons exhibit dye integration throughout their architecture, suggesting that the EVs have fused and/or loaded their content into these cells. Soluble Bodipy dye at the concentration used to label the vesicles does not label neurons, ruling out the possibility that the observed neuronal labeling is caused by residual, non-vesicular dye. Lys=lysate; CD=cell debris.

**Supplementary Figure 2.**
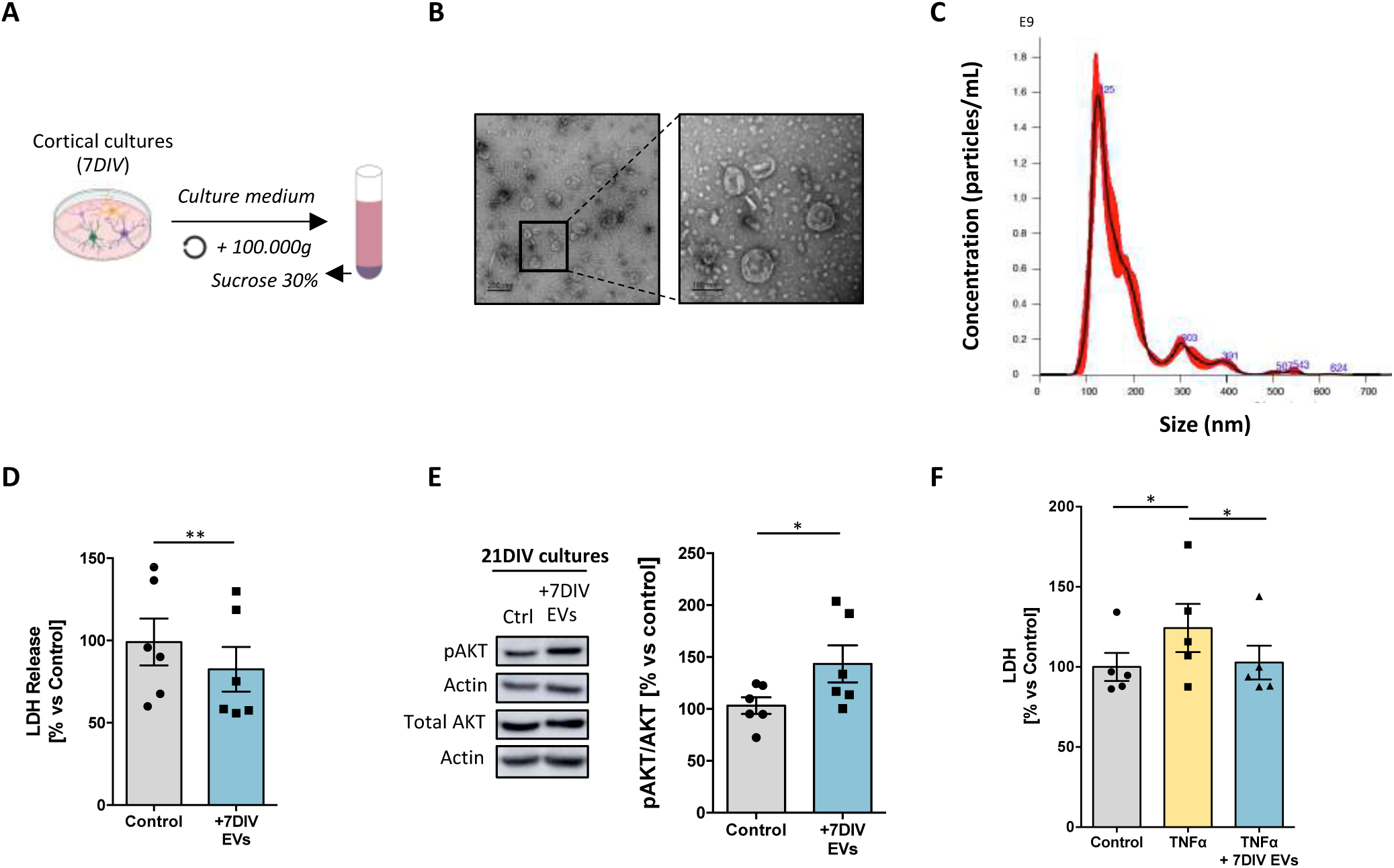
Small extracellular vesicles derived from young neuronal cultures (7DIV EVs) have a beneficial effect on aged 21 DIV neuronal cultures. **(A)** Schematic representation of vesicle preparation from cultured cells. **(B)** Representative transmission electron microscopy (TEM) image of vesicles obtained from the medium of mouse neuronal cultures by centrifugation at 100,000 x g. **(C)** Representative size distribution of the vesicles present in the conditioned medium of mouse neuronal cultures and analyzed by Nanoparticle Tracking Analysis (NTA). The profile shows that most of the vesicles have a size between 50 and 200 nm. **(D)** Absorbance measurement of LDH release into the conditioned medium of 21DIV cultures treated (+EVs 7DIV) or not (Control) with 7DIV EVs at 1 x 10^9^ particles/mL for 24 h (n = 6; **p < 0.01; Student’s t-test). **(E)** Western blot analysis of the ratio of the levels of activated AKT protein (pAKT) to total protein (AKT) in 21DIV cultures treated or not with 7DIV EVs (n = 6; *p < 0.05; Student’s t-test). Actin was used as a loading control. **(F)** Absorbance measurement of LDH release into the conditioned medium of 14DIV cultures treated or not with TNFα (50 ng/mL) and in the presence or absence of 7DIV EVs (n = 5; *p < 0.05; one-way ANOVA followed by Tukeýs post hoc correction). All data represent the mean ± SEM as a percentage of the control (untreated) values.

**Supplementary Figure 3.**
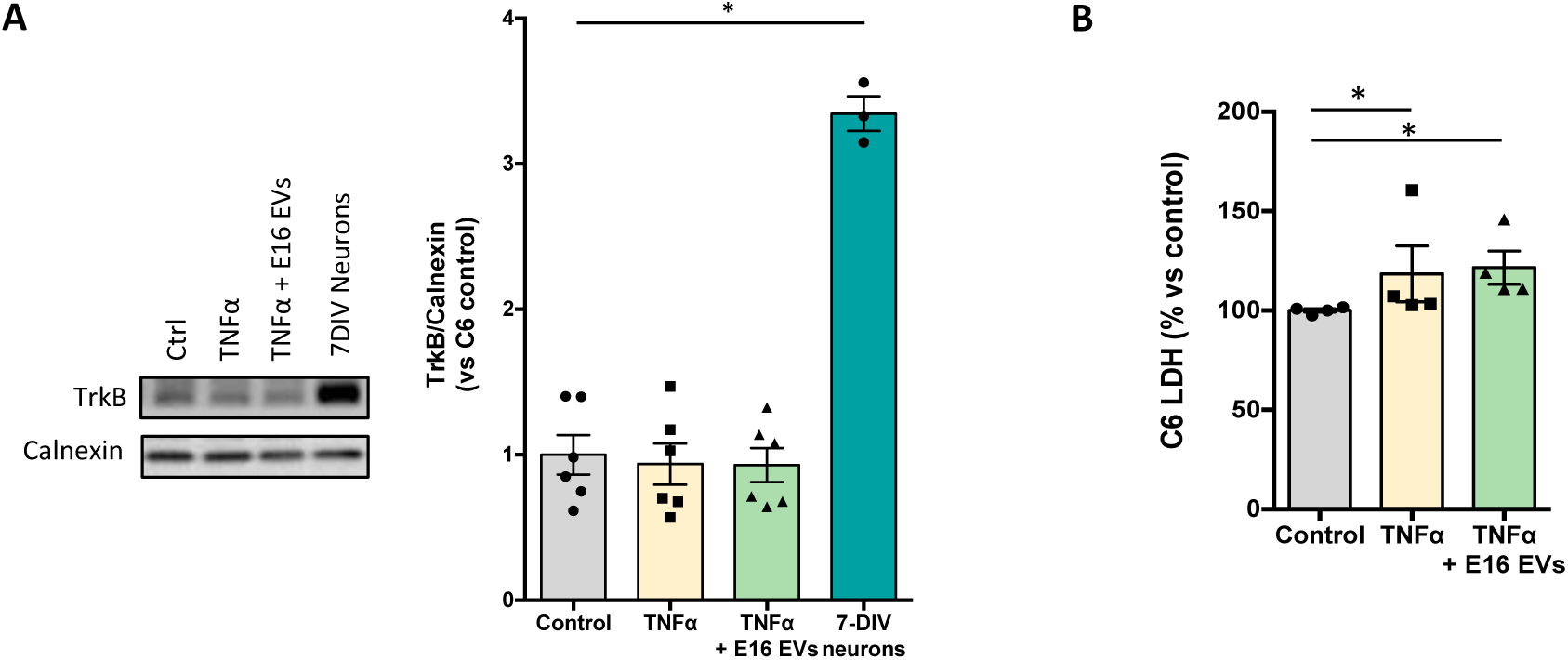
E16 EVs fail to diminish TNF-α-induced LDH release in C6 glial cells. **(A)** Western blot analysis of total TrkB protein in C6 glial cells treated or not with TNFα (50 ng/mL) in the presence or absence of E16 EVs. TrkB activation levels in 7 DIV neurons were also compared with those in C6 cells. Calnexin was used as a loading control (n = 6; *p < 0.05; one-way ANOVA followed by Tukeýs post hoc correction). **(B)** Absorbance measurement of LDH release into the conditioned medium of C6 glial cells treated or not with TNFα (50 ng/mL) in the presence or absence of E16 EVs (n = 4; *p < 0.05; one-way ANOVA followed by Tukeýs post hoc correction). All data represent the mean ± SEM as a percentage of the control (untreated) values.

## AUTHORS CONTRIBUTION

R.G-R. performed the majority of the experiments, collected data and conducted the statistical analysis; M.G-V. collaborated in the *in vivo* experiments; M.C-C. performed experiments using the C6 cell line; S.G.F. carried out the bioinformatic analysis of proteomics data; I.C.P. and C.C. advised on and contributed to the isolation of EVs by size-exclusion chromatography; C.G.D. and F.X.G. designed the study and conceptualized the research; C.G.D wrote the initial draft; C.G.D, E.P, F.G.X and R.G-R. reviewed and edited the manuscript. All authors have read and approved the final manuscript.

## FUNDING

This work was supported by Agencia Estatal de Investigación (MICIU/AEI/10.13039/501100011033) through grants PID2019-104389RB-I00 (ERDF/EU) to C.G.D, including the award of a predoctoral fellowship to R.G-R. for the training of research personnel (FPI) and PID2022-138334OB-I00 (ERDF/EU) to C.G.D and F.G.R.

## ACKNOWLEDGMENTS

We thank the different scientific services of the Centro de Biología Molecular Severo Ochoa (CBM): the Electron Microscopy Service for sample staining and processing; the Confocal Microscopy Service for their assistance with image analysis; and the Animal Facility for the care of the experimental animals. We also thank the group of Jesús Vázquez at the CNIC, particularly Emilio Camafeita, for carrying out the proteomics experiments as well as their statistical analysis. We thank Miguel Ángel Marchena Fernández and Fernando de Castro from Cajal Institute for their assistance in the design of the retinal experiments. We thank Dr. Kenneth McCreat for editorial assistance with English language, grammar, and style.

## Notes

### Competing Interest Statement

The authors have declared no competing interest.

